# A new Persister Enrichment Model in *Mycobacterium tuberculosis* links multidrug persistence to drug sequencing and maltokinase-dependent carbon distribution

**DOI:** 10.64898/2026.05.28.728405

**Authors:** Michael W. Shultis, Claire V. Mulholland, Jinhua Cui, Mei Chen, Samantha Johnson, Christopher Hiner, Michael Berney

## Abstract

Prolonged combination chemotherapy is required to cure Tuberculosis, in part because *Mycobacterium tuberculosis* (*Mtb*) forms antibiotic-tolerant persister cells that survive exposure to bactericidal drugs. Experimental systems to study multidrug persistence in *Mtb* remain limited. Here, we describe a rapid and tractable Persister Enrichment Model (PEM) in which removal of glycerol markedly increases the frequency of multidrug-tolerant *Mtb* cells that survive exposure to high concentrations of bactericidal antibiotics with distinct mechanisms of action. Persister enrichment in PEM depends on carbon source identity rather than carbon abundance and enables direct interrogation at the transition from active growth into multidrug tolerance. Using PEM, we show that the temporal order of isoniazid (INH) and rifampicin (RIF) exposure shapes the magnitude of multidrug persistence. A transposon sequencing screen under combination drug pressure identified maltokinase (*mak*) as a major determinant of multidrug persistence. Deletion of *mak* enhanced susceptibility to INH+RIF in PEM and impaired bacterial survival during the persistent phase of infection in mice. Metabolic analyses reveal that loss of *mak* destabilizes carbon distribution during co-catabolism of glycolytic carbon sources, leading to defective growth and antibiotic persistence. This defect is ameliorated by inactivating mutations in glucose-1-phosphate adenylyltransferase (*glgC*), which alleviate the buildup of ADP-glucose. Together, these findings establish PEM as a practical platform to study multidrug persistence, reveal that drug sequencing can influence entry into multidrug tolerance, and identify maltokinase-dependent carbon distribution as a metabolic vulnerability that constrains *Mtb* survival.

**Importance:** Tuberculosis treatment requires prolonged multidrug chemotherapy in part because *Mycobacterium tuberculosis* forms antibiotic-tolerant persister cells that survive bactericidal drug exposure. However, experimental systems to study multidrug persistence remain limited, and the metabolic mechanisms underlying this phenotype are poorly understood. Here, we establish a rapid and experimentally tractable Persister Enrichment Model (PEM) that enriches multidrug-tolerant *M. tuberculosis* populations by removing glycerol from culture medium. Using this model, we demonstrate that the order of antibiotic exposure influences multidrug persistence both *in vitro* and *in vivo* and identify maltokinase-dependent carbon distribution as a critical determinant of persistence and long-term bacterial survival. These findings provide a new platform for studying multidrug tolerance and reveal carbon buffering pathways as potential therapeutic vulnerabilities that could help shorten tuberculosis treatment.

## Introduction

Tuberculosis (TB) has re-emerged as the leading cause of death from a single infectious agent worldwide, accounting for approximately 1.23 million deaths annually^1^. TB chemotherapy is arduous and frequently associated with severe adverse effects, including ocular toxicity, peripheral neuropathy, gastrointestinal distress, fever, and profound fatigue^2^. Notably, clinical symptoms often resolve months before completion of the required drug regimen, leaving patients exposed to treatment-related toxicity in the absence of overt disease^3^. Patient-centered studies and meta-analyses have highlighted the substantial burden imposed by TB therapy, with some patients discontinuing treatment prematurely or continuing despite significant detriments to quality of life^4^. These challenges emphasize the urgent need to identify strategies that reduce treatment duration and toxicity while maintaining efficacy.

A major barrier to shortening TB therapy is antibiotic persistence, a phenotypic state in which a subpopulation of bacteria survives exposure to bactericidal concentrations of antibiotics without acquiring heritable resistance mutations^5,6^. The requirement for extended multidrug chemotherapy in TB has long been attributed to the presence of persistent bacilli that survive initial treatment, necessitating months of continued therapy^7–9^. Prior work has identified both stochastic and stress-induced mechanisms of persister formation in *Escherichia coli*, with diverse underlying processes including metabolic remodeling and toxin-antitoxin systems^10,11^. In mycobacteria, intrinsic cellular heterogeneity arising from asymmetric growth and differential metabolic states has been shown to influence antibiotic susceptibility, suggesting that persister populations are not uniform^12,13^. Consistent with this, evidence indicates that persistence is often drug-specific, with distinct subpopulations exhibiting incomplete and asymmetric cross-tolerance between antibiotics^14–17^. Notably, rifampicin and pyrazinamide exhibit enhanced activity against persistent populations, relative to both other antibiotics and their activity against actively growing cells, further highlighting functional differences among persister states^18^. However, whether persisters selected by one drug overlap with those surviving multidrug therapy remains poorly understood^5^. Because TB treatment relies exclusively on combination chemotherapy, multidrug persisters represent a critical but poorly understood obstacle to sterilizing cure. Persistence to frontline drugs, including isoniazid (INH), rifampicin (RIF), ethambutol (EMB), and pyrazinamide (PZA), is therefore of particular concern.

Antibiotic persistence in *Mycobacterium tuberculosis* (*Mtb*) is typically interrogated using models that induce a non-replicative state in the bacteria, such as PBS starvation^19^, stationary phase culture^20^, or hypoxia^21^. Setup for these models can require up to six weeks of preparation prior to the administration of drug treatment. Here, we introduce a defined *in vitro* model to investigate multidrug persistence in *Mtb*. The persistent population generated by this model exhibits key features consistent with established definitions of persistence, such as bimodal killing and regrowth after removal of antibiotics^22^. Using this system, we demonstrate that bacterial persistence is strongly influenced by the order of drug administration in combination therapies, consistent with triggered persistence^22^. Applying the model to the frontline INH+RIF regimen, we identify maltokinase (*mak*) as a key modulator of multidrug persistence *in vitro* and as a determinant of bacterial survival during the persistent phase of infection *in vivo*. Together, these findings establish a tractable framework for dissecting multidrug persistence and reveal a metabolic vulnerability that may be exploited to improve TB treatment outcomes.

## Results

### Glycerol removal establishes a rapid Persister Enrichment Model (PEM) for multidrug persistence

We observed that the removal of glycerol from standard 7H9 broth medium (7H9/OADC/tyloxapol containing 2 mg/mL dextrose, hereafter referred to as ‘Dex2’) markedly enriched the drug-tolerant survivor population after only eight days of culture prior to antibiotic exposure (Figure 1A-F, Supplemental Figure 1). Bacterial cultures were grown to mid-log phase in the presence or absence of glycerol and then treated with antimycobacterial agents at 50-100× their MIC at ∼10^8^ CFU/mL. We evaluated three frontline drugs (INH, RIF, and EMB), alone or in combination, to model a multidrug persister population most closely representative of patients receiving clinical treatment. All treatments, excluding RIF monotherapy, resulted in a 2-4 log_10_ increase in surviving bacteria, indicating enrichment of drug-tolerant populations across multiple antibiotic classes (Figure 1A-E). This phenotype was observed in both avirulent *Mtb* mc^2^6230 and virulent *Mtb* H37Rv (Supplemental Figure 1A). Unless otherwise noted, subsequent experiments were performed using the mc^2^6230 strain. We also tested additional drugs with different mechanisms of action, such as the cell wall inhibitor D-cycloserine (DCS), the protein synthesis inhibitor linezolid (LNZ), and the DNA gyrase inhibitor ofloxacin (OFX), all of which showed enrichment of the persister population in the absence of glycerol (Supplemental Figure 1B).

**Figure 1.**
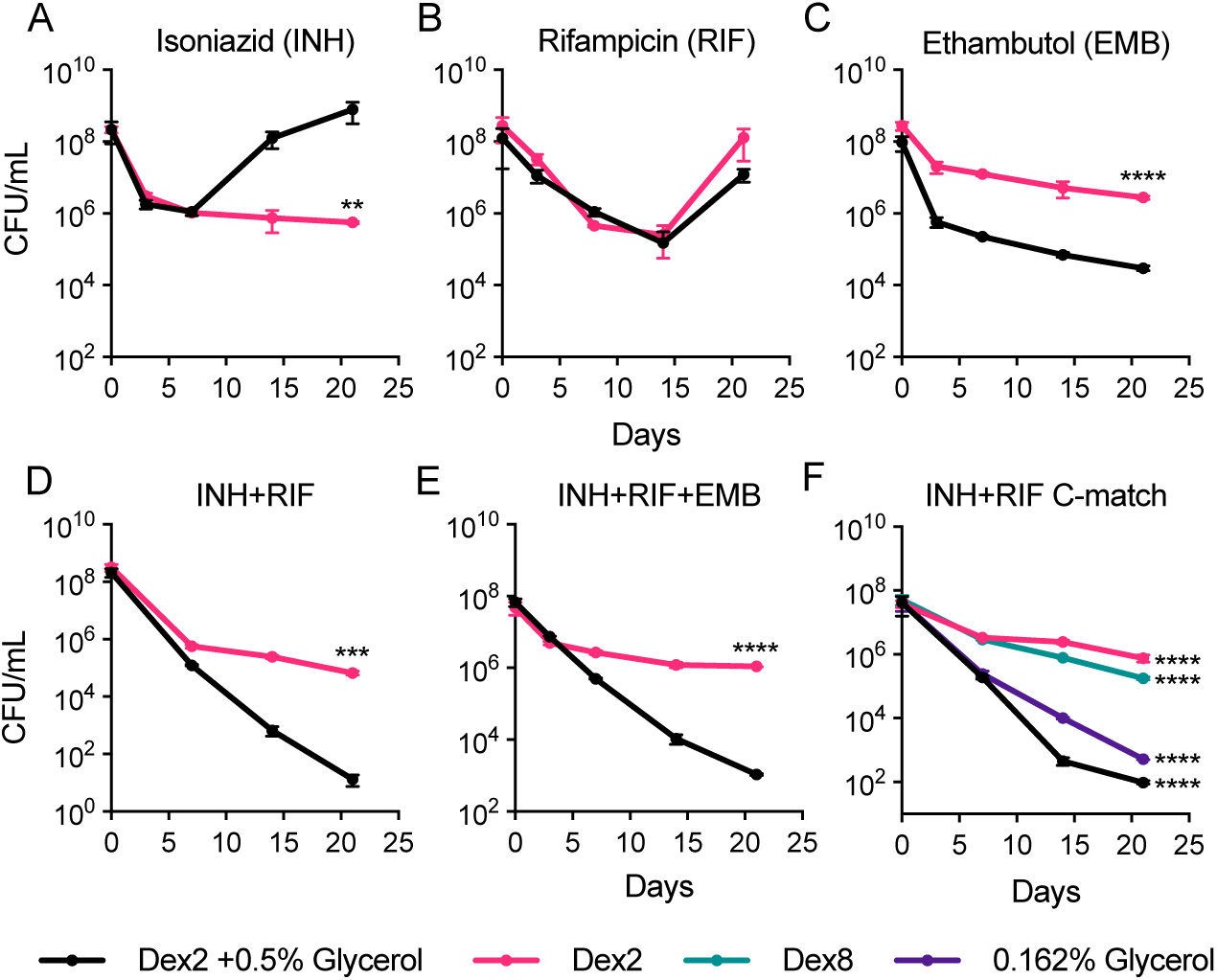
Removal of glycerol from culture media enhances *Mtb* survival to drug combinations due to altered glycerol-dependent physiology. **(A-E)** *Mtb* was grown to mid-log phase in the presence or absence of glycerol, then challenged with different combinations of INH (36 µM), RIF (3 µM), and EMB (490 µM) in Dex2 media with glycerol (black) or without glycerol (pink). Data represent mean ± SD (n = 3). Statistical significance was determined by Student’s t-test comparing +glycerol and *-*glycerol at the final timepoint (**p <0.01, ***p <0.001, ****p <0.0001). **(F)** *Mtb* was grown to mid-log phase in Dex2 media with glycerol (black), Dex2 without glycerol (pink), Dex8 (green; dextrose only, carbon-matched to Dex2 + 0.5% glycerol), and 0.162% glycerol (purple; glycerol only, carbon-matched to Dex2). Cultures were treated INH (36 µM) + RIF (3 µM) (I). Data represent mean ± SD (n = 3). Statistical significance was determined by one-way ANOVA with Tukey’s multiple comparisons test at the final timepoint. Asterisks denote Tukey-adjusted pairwise comparisons; asterisks indicate that all comparisons with the tested group were significant, while brackets indicate specific pairwise comparisons between groups (****p <0.0001). Data are representative of at least two independent experiments. **Dex2** = 7H9/OADC/tyloxapol with a final concentration of 2 mg/mL dextrose, **Dex8** = 7H9/OADC/tyloxapol with a final concentration of 8 mg/mL dextrose, **0.162% Glycerol** = 7H9/OAC/tyloxapol/glycerol with a final concentration of 0.162% glycerol.

To determine whether the PEM phenotype reflected reduced carbon availability and define the role of glycerol in persister formation, we established media conditions that decouple carbon quantity from carbon source identity (Figure 1F). This included glycerol-only medium (0.162% glycerol), matching the total carbon content of the Dex2 PEM condition, and a dextrose-only medium (Dex8), matching the total carbon content of Dex2 supplemented with 0.5% glycerol. This design enabled independent assessment of the effects of carbon source identity versus carbon abundance on persister formation under INH+RIF co-treatment (Figure 1F). Even low concentrations of glycerol abolished persister enrichment, whereas high dextrose retained it (Figure 1F). Because all media contained sodium oleate, we also verified that oleate did not influence the persister enrichment phenotype (Supplemental Figure 2). These results indicate that glycerol exerts a specific metabolic effect not recapitulated by excess sugar alone. Consistent with prior studies linking glycerol metabolism to altered antibiotic susceptibility^23^, suppression of the PEM phenotype is closely associated with glycerol-dependent physiology. In contrast, glycerol-free conditions consistently supported persister enrichment (Figure 1, Supplemental Figure 1).

### PEM expands the experimental window for studying isoniazid persisters

INH is a cornerstone of therapy for drug-susceptible *Mtb*. Although INH persistence has been studied before^24^, commonly used models provide a narrow window in which only a small transient persister population can be observed before resistant mutants rapidly dominate the culture^24^. PEM overcomes this limitation as cultures are not rapidly overtaken by resistant bacteria and the persister population size is substantially increased (Figure 1A). Longitudinal analysis of INH kill curves showed that glycerol-containing cultures were predominantly resistant by day seven and fully resistant by day 14, whereas glycerol-free cultures remained largely non-resistant (<5% resistant on day 7, ∼20% on day 14), with complete resistance emerging only by day 35 (Supplemental Figure 3). The predominant resistance mutations identified under both conditions were *katG* frameshift mutations (Supplemental Table 1), the gene most commonly associated with INH resistance in clinical isolates^25^. Together, these data establish PEM as a system that extends both the duration and scale of the INH persister population prior to resistance takeover.

### Drug exposure sequence shapes multidrug persistence *in vitro* and *in vivo*

Having established a robust INH persister population, we next used PEM to examine cross-tolerance and the impact of drug scheduling on multidrug persistence. Cultures were treated with INH for seven days, followed by the addition of a second drug (Figure 2A). INH persisters remained tolerant to additional INH but showed reduced tolerance to EMB and RIF (Figure 2A). Sequential treatment with INH followed by RIF resulted in greater bacterial killing than simultaneous INH+RIF treatment (Figure 2B). In contrast, initiating treatment with RIF prior to INH reduced the efficacy of the combination over the same time frame (Figure 2B). These results indicate that *Mtb* persister populations undergo drug-dependent changes in susceptibility to subsequent treatments, consistent with triggered persistence^22^ and prior models of multidrug tolerance^5,26^.

**Figure 2.**
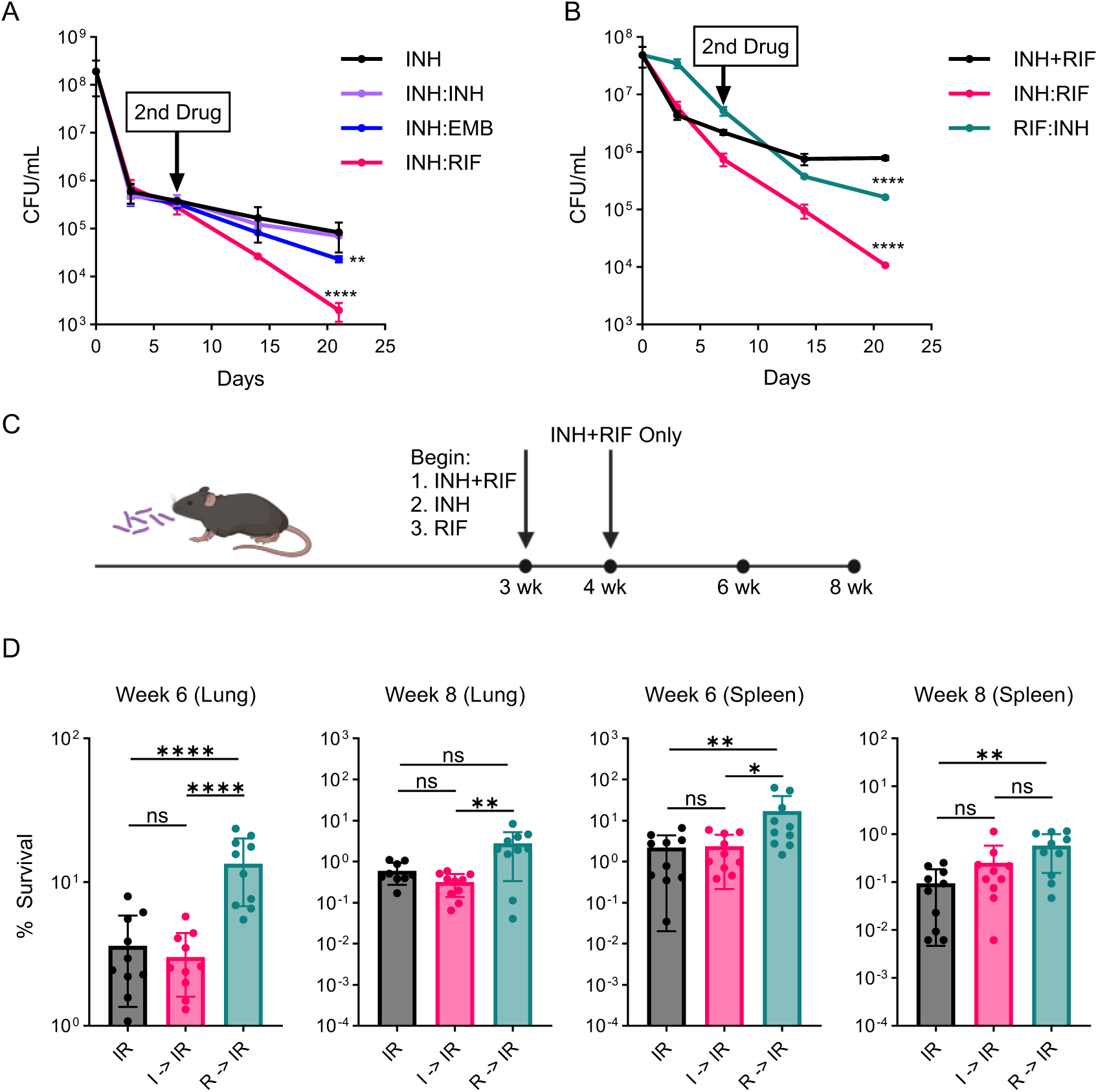
Drug scheduling influences *Mtb* persister formation and drug tolerance in PEM and mice. **(A)** PEM cultures were treated with INH (36 µM) for 7 days, followed by the addition of additional INH (36 µM), EMB (490 µM), or RIF (3 µM). Data represent mean ± SD (n = 3). Statistical significance was determined by one-way ANOVA with Tukey’s multiple comparisons test comparing all conditions to INH at the final timepoint (**p <0.01, ****p <0.0001). **(B)** PEM cultures were treated with simultaneous INH+RIF, sequential INH followed by RIF (INH:RIF), or sequential RIF followed by INH (RIF:INH), with a 7-day interval between sequential drug additions. Data represent mean ± SD (n = 3). Statistical significance was determined by one-way ANOVA with Tukey’s multiple comparisons test comparing all conditions at the final timepoint (****p <0.0001). **(C-D)** C57BL/6 mice were aerosol-infected with H37Rv and treated with INH (100 mg/L) and RIF (40 mg/L) administered *ad libitum* in drinking water starting 3 weeks post-infection. Mice received either combination therapy from treatment initiation (IR, black) or one week of INH or RIF monotherapy followed by combination therapy (I→IR, Pink; R→IR, green). Data represent mean ± SD from two independent experiments (n = 10). Statistical significance was determined by one-way ANOVA with Tukey’s multiple comparisons test at each timepoint within each organ (*p <0.05, **p <0.01, ****p <0.0001). Apart from 2A, data are representative of at least two independent experiments.

To assess whether these sequence-dependent effects extend *in vivo,* we infected C57BL/6 mice via aerosol with *Mtb* H37Rv. After three weeks of infection, mice were treated with INH, RIF, or INH+RIF administered *ad libitum* in drinking water. Animals receiving INH or RIF monotherapy were switched to combination therapy after one week, and bacterial burden was measured after three and five weeks of treatment (Figure 2C). Sequential administration of RIF followed by INH+RIF resulted in reduced bacterial killing in both the lungs and spleens compared to simultaneous combination therapy (Figure 2D), consistent with our *in vitro* findings (Figure 2B). In contrast, sequential administration of INH followed by INH+RIF did not significantly alter bacterial burden (Figure 2D). Together, these findings demonstrate that the order of INH and RIF exposure influences treatment efficacy *in vivo*.

### Maltokinase modulates multidrug persistence

Having established PEM as a robust system for generating and studying multidrug persistent *Mtb* and revealing that INH+RIF combination treatment results in a more substantial persister population, we next sought to identify genetic determinants of multidrug persistence. To accomplish this, we performed Tn-Seq under combination drug pressure. A saturated transposon mutant pool was treated with EMB alone, INH+RIF, or INH+RIF+EMB under glycerol-containing and glycerol-free conditions (Figure 3A). To identify genes conditionally required for multidrug persistence, data from all drug treatments were pooled for analysis, and the glycerol-containing and glycerol-free conditions were compared. This analysis identified 322 genes with significant fitness effects, of which 231 were more essential for survival in the absence of glycerol (Figure 3B, Supplemental Table 2).

**Figure 3.**
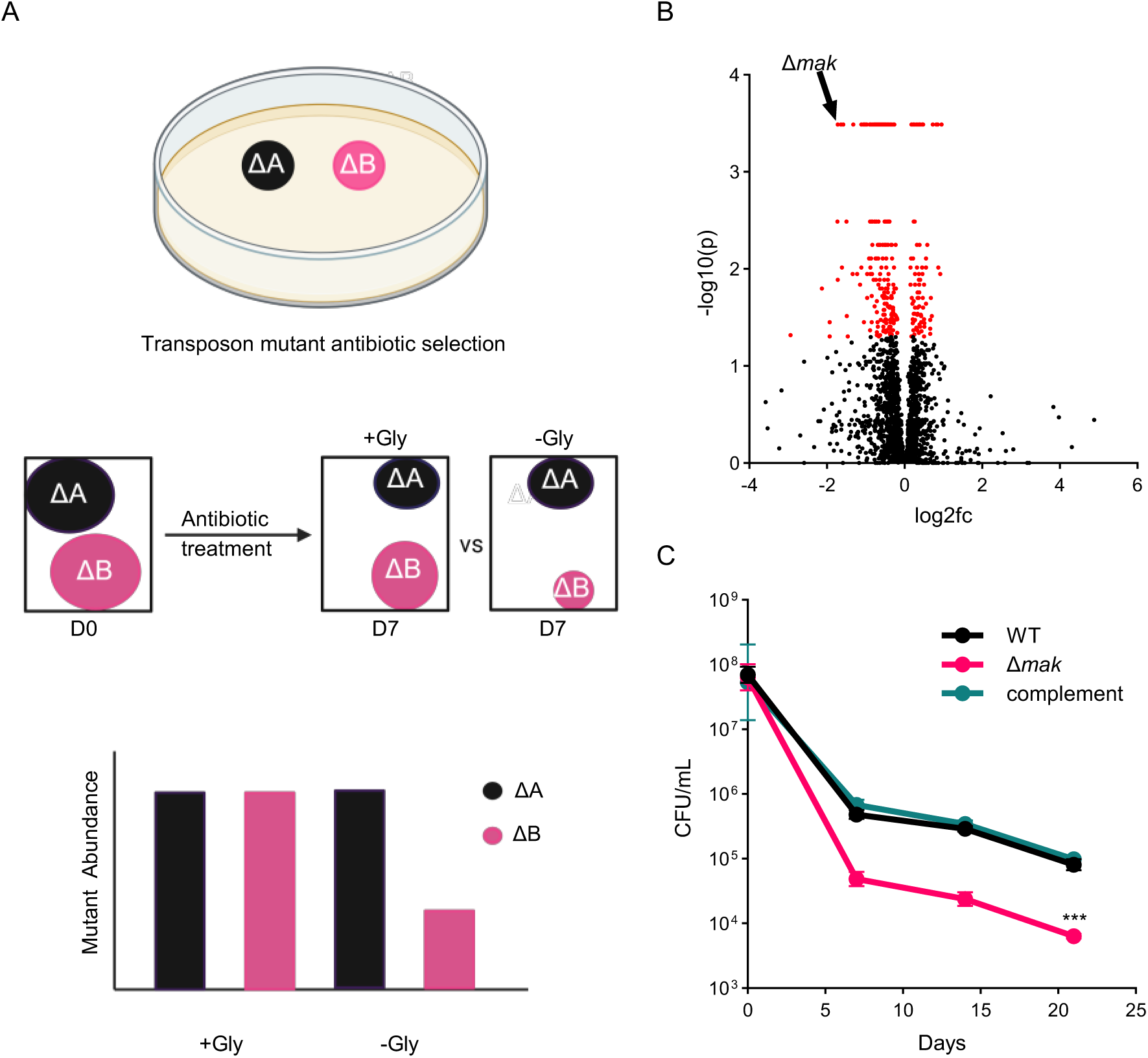
Tn-Seq reveals maltokinase as a multidrug persistence factor. **(A)** A schematic representation of the transposon mutagenesis workflow. First, conditionally essential mutants were isolated using antibiotic selection media. This pool of mutants was then treated with antibiotics (EMB, INH+RIF, or INH+RIF+EMB) for 7 days in the presence and absence of glycerol. Some mutants, such as ΔA, were identically represented in both conditions, while others, such as ΔB, were differentially represented, indicating genes that influence persister enrichment. **(B)** A volcano plot representing the screen of transposon mutants tested after drug treatment in the presence of glycerol versus the absence of glycerol. A negative log_2_ fold change represents an underrepresented mutant in our PEM compared to Dex2 media with glycerol. **(C)** Wild-type, Δ*mak,* and the complemented strain were treated with INH+RIF in our PEM, validating Δ*mak* as a multidrug persister gene. Data represent mean ± SD (n = 3). Statistical significance was determined by Student’s t-test comparing wild-type and Δ*mak* at the final timepoint. (***p <0.001). Data are representative of at least two independent experiments. **Dex2** = 7H9/OADC/tyloxapol with a final concentration of 2 mg/mL dextrose. INH (36 µM), RIF (3 µM), and EMB (490 µM).

Among the most strongly depleted genes was the maltokinase-encoding gene *mak*, which exhibited a 3.3-fold decrease in persistence in the absence of glycerol (Figure 3B). This gene stood out for several reasons: (1) it was consistently underrepresented in the surviving population under drug pressure, (2) it encodes a key enzyme in the TreS-Mak-GlgE α-glucan biosynthesis pathway that has previously been implicated in antibiotic persistence^27^, and (3) the maltokinase enzyme (Mak) operates at the interface of carbon storage and sugar metabolism, representing a compelling candidate to study the metabolic basis of multidrug persistence. To validate its role in multidrug persistence, we generated a targeted deletion mutant by specialized transduction^28^. In PEM, *Mtb* Δ*mak* showed significantly reduced survival during INH+RIF treatment, and complementation under the native promoter restored survival to wild-type levels (Figure 3C), establishing maltokinase as a determinant of multidrug persistence. This effect was recapitulated in a virulent H37Rv background (Supplemental Figure 4).

Maltokinase, also known as “Pep2”, functions in the TreS-Mak-GlgE glycogen biosynthesis pathway^29–31^, catalyzing the conversion of maltose to α-maltose-1-phosphate, which is subsequently utilized by GlgE to generate the branching sugar chains of the carbon storage molecule glycogen (Figure 4A). GlgE is considered essential in *Mtb* due to the toxic accumulation of α-maltose-1-phosphate when GlgE is disrupted^29^. The upstream enzyme, TreS, was also identified in our screen and has previously been implicated in antibiotic persistence through a mechanism described as the “trehalose catalytic shift”^27^. This mechanism proposes that conversion of trehalose to maltose enables its utilization in central carbon metabolism^27^. However, because *Mtb* Δ*mak* retains this capacity, the observed persistence defect suggests that maltokinase contributes to persistence through downstream regulation of carbon distribution rather than trehalose entry per se.

**Figure 4.**
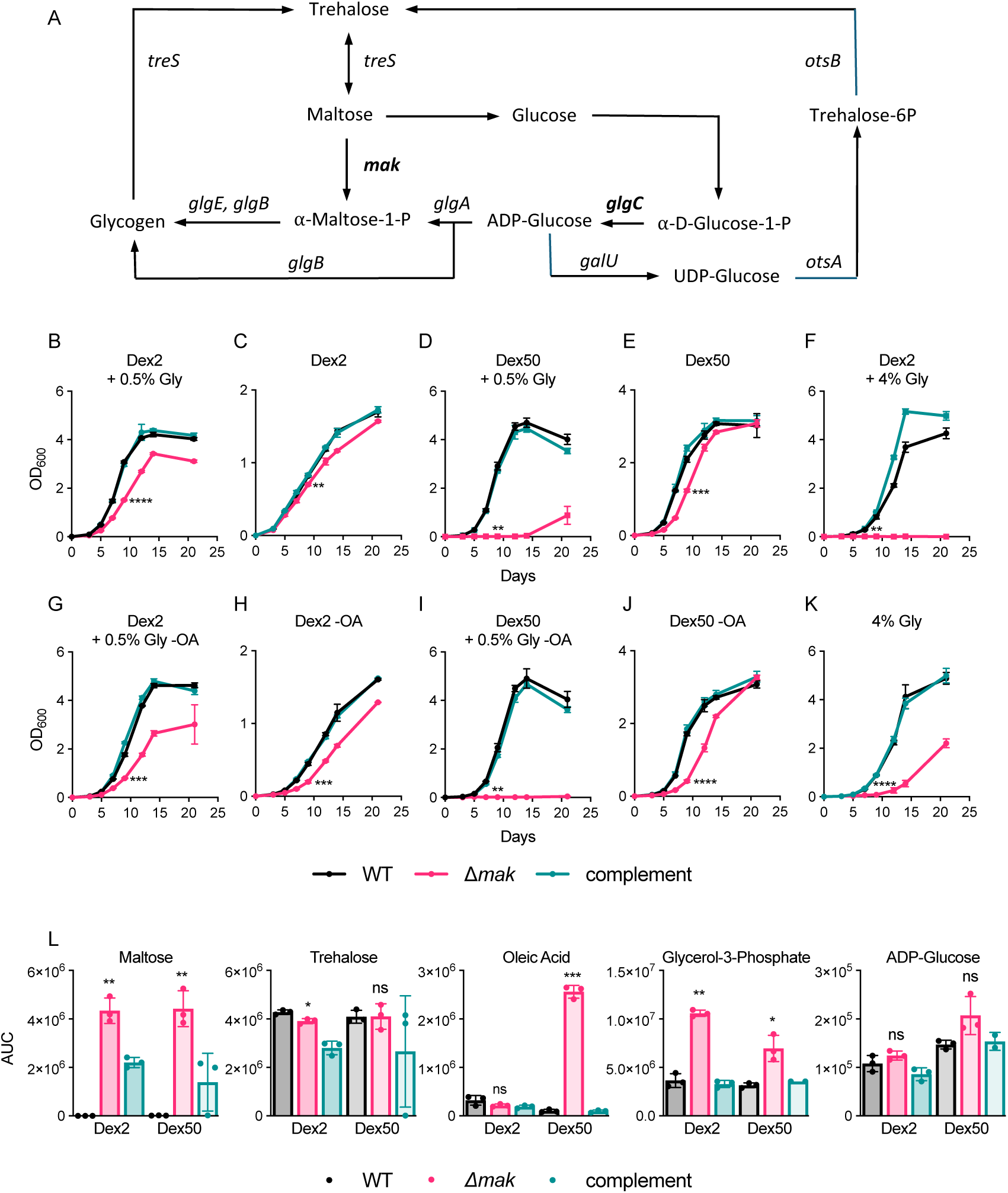
*Δmak* exhibits metabolic vulnerability during co-catabolism of glycerol and dextrose, resulting in oleate and ADP-glucose accumulation. **(A)** A schematic representation of relevant intermediates and enzymes in glycogen synthesis. **(B-K)** Growth curves of wild-type (black), Δ*mak* (pink), and complement (green) grown under various carbon source combinations as indicated. Data represent mean ± SD (n = 3). Statistical significance was determined using a Student’s t-test comparing wild-type and mutant at day 9 (*p <0.05, **p <0.01, ***p <0.001, ****p <0.0001). **(L)** Abundances of significantly altered metabolites involved in glycogen synthesis, along with oleate, identified from a semi-targeted metabolomic analysis of ∼190 individual metabolites comparing the wild-type and Δ*mak* strains under standard Dex2 media conditions and 50 mg/mL dextrose +0.5% Glycerol (Dex50) (Supplemental Table 3). Oleate was the only metabolite exhibiting a log_2_fc >10 in Dex50 conditions. Data are shown as area under the curve (AUC) and represent mean ± SD (n = 3). Statistical significance was determined by multiple Student’s t-tests adjusted using the Benjamini-Hochberg procedure (q-values) (*q <0.05, **q <0.01, ***q <0.001). Data are representative of at least two independent experiments. **Dex2** = 7H9/OADC/tyloxapol with a final concentration of 2 mg/mL dextrose, **Dex50** = 7H9/OADC/tyloxapol with a final concentration of 50 mg/mL dextrose, **4% Gly** = 7H9/OAC /tyloxapol/glycerol with a final concentration of 4% glycerol, **+0.5% Gly** = Supplementation with 0.5% glycerol, +**4% Gly** = Supplementation with 4% glycerol, **-OA** = Oleic acid removed.

Notably, the persistence defect of Δ*mak* was specific to PEM conditions, as the mutant exhibited survival comparable to the wild-type in the presence of glycerol (Supplemental Figure 5). While both Δ*mak* and glycerol independently reduced survival, glycerol exposure produced a stronger effect and did not further exacerbate the Δ*mak* phenotype, suggesting overlapping metabolic constraints. These findings are consistent with a model in which loss of maltokinase perturbs carbon distribution^32^ and partially recapitulates a glycerol-associated metabolic stress state, in line with prior work linking glycerol metabolism to altered drug tolerance^26,33^.

### *Mtb* Δ*mak* reveals a synthetic toxicity during co-catabolism of glycolytic carbon sources

In addition to its persistence phenotype, Δ*mak* displayed a growth defect in Dex2 medium in the presence of glycerol, which was restored by complementation (Figure 4B). This defect was also observed in an H37Rv background (Supplemental Figure 6). This defect was less severe in glycerol-free medium (Figure 4C). Supplementation with additional dextrose in glycerol-containing medium further exacerbated the growth defect (Figure 4D), consistent with a synthetic toxicity under co-catabolic carbon conditions. Supplementation with additional dextrose in the absence of glycerol did not exhibit the same toxicity, indicating glycerol dependence (Figure 4E).

To investigate the metabolic basis of this phenotype, we performed metabolic profiling of wild-type, Δ*mak,* and complemented strains using established protocols^34,35^. Semi-targeted analysis of approximately 190 metabolites revealed widespread metabolic perturbations in Δ*mak* under both standard and high-dextrose conditions (Supplemental Table 3). As expected, maltose accumulated in Δ*mak*, whereas trehalose showed minimal change (Figure 4L).

Oleic acid was uniquely increased in high-dextrose media, while glycerol-3-phosphate was elevated in Δ*mak* under both conditions (Figure 4L). Oleic acid, after conversion to oleolyl-CoA is a major building block for triacylglycerides (TAG)^36^. Because *Mtb* accumulates triacylglycerides (TAGs) during arrested growth on fatty acids^36^ we tested whether oleic acid abundance influences the Δ*mak* growth defect. Removal of oleate from culture media exacerbated the defect across all conditions, with the most severe impairment observed under Dex50 supplemented with 0.5% glycerol (Figure 4G-J). These data suggest that oleate partially mitigates glycerol-associated toxicity, potentially by facilitating sequestration of glycerol via TAG synthesis.

To compare the influence of glycerol to that of dextrose, we increased glycerol concentrations to match those of Dex50 conditions. Increasing glycerol from 0.5% to 4% impaired the growth of Δ*mak* relative to wild-type and complemented strains (Figure 4K) with an even more pronounced defect in Dex2 +4% glycerol (Figure 4F). Together, these findings indicate that the most toxic defect arises during the co-catabolism of sugar and glycerol, with defective growth being influenced more by glycerol than dextrose, regardless of concentration.

### Inactivating mutations in *glgC* alleviate growth toxicity in Δ*mak* by preventing ADP-glucose accumulation

During growth in Dex50 supplemented with 0.5% glycerol, Δ*mak* cultures showed prolonged growth arrest followed by eventual outgrowth, suggesting the acquisition of suppressor mutations (Figure 4D). Individual colonies were isolated from the outgrowth population and analyzed by whole-genome sequencing, revealing mutations in two genes (Supplemental Table 4). The most frequent was insertion of an IS*6110* mobile element into *glgC*, which encodes glucose-1-phosphate adenylyltransferase (GlgC), an enzyme in the alternative glycogen biosynthesis pathway (Figure 4A). GlgC catalyzes the conversion of α-D-glucose-1-P to ADP-glucose, a precursor that can be incorporated into glycogen or diverted into α-maltose-1-P by GlgA^37^ (Figure 4A). The second mutation was a non-synonymous G62D substitution in *sugI*, a probable sugar transporter^38,39^. Both mutations restored growth in Dex50 +0.5% glycerol media (Figure 5A). However, only Δ*mak glgC*::IS*6110* restored growth in Dex2 +4% glycerol media (Figure 5B-C). This is consistent with the proposed function of SugI, as limiting sugar uptake alleviated toxicity under high dextrose conditions but did not mitigate glycerol-associated toxicity. In contrast, *glgC* disruption restored growth under both conditions, implicating GlgC activity in the glycerol-associated Δ*mak* growth defect.

**Figure 5.**
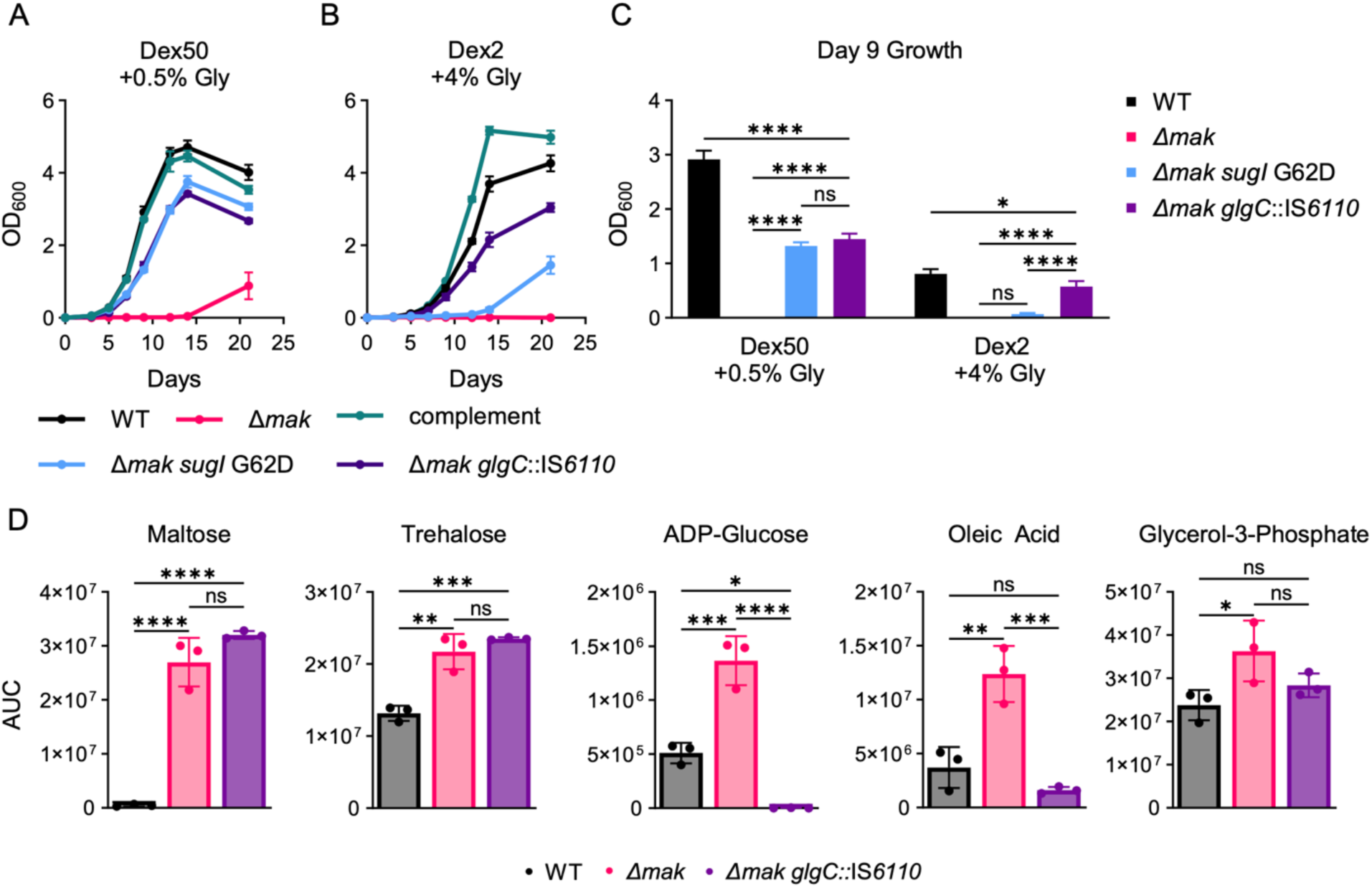
ADP-Glucose toxicity is alleviated by suppressor mutations in *glgC.* **(A-B)** Growth curves of wild-type (black), Δ*mak* (pink), complement (green), Δ*mak glgC*::IS*6110* (purple), and Δ*mak sugI* G62D (blue) grown under two high-carbon conditions. Data represent mean ± SD (n = 3). **(C)** Growth on day 9 from the growth curves (A-B) showing statistical significance was determined by two-way ANOVA. For clarity, statistical comparisons are shown at a single representative timepoint, selected to reflect the overall experimental trend. (*p <0.05, ****p <0.0001). **(D)** LC-MS analysis of five metabolites identified in the prior semi-targeted analysis was performed in wild-type (black), Δ*mak* (pink), and Δ*mak glgC*::IS*6110* (purple) strains in Dex50 +0.5% glycerol media. Data represent mean ± SD (n = 3). Statistical significance was determined by one-way ANOVA of each metabolite with Tukey’s multiple comparisons test. (*p <0.05, **p <0.01, ***p <0.001, ****p <0.0001). Data are representative of at least two independent experiments. **Dex2** = 7H9/OADC/tyloxapol with a final concentration of 2 mg/mL dextrose, **Dex50** = 7H9/OADC/tyloxapol with a final concentration of 50 mg/mL dextrose, **+0.5% Gly** = Supplementation with 0.5% glycerol, +**4% Gly** = Supplementation with 4% glycerol.

To directly assess the metabolic impact of *glgC* disruption, we measured the abundance of trehalose, maltose, glycerol-3-phosphate, oleic acid, and ADP-glucose in wild-type, Δ*mak*, and Δ*mak glgC*::IS*6110* strains in Dex50 + 0.5% glycerol media (Figure 5D, Supplemental Table 5). ADP-glucose was markedly reduced in Δ*mak glgC*::IS*6110*, consistent with the loss of GlgC activity (Figure 5D). This was accompanied by the normalization of oleate levels and partial normalization of glycerol-3-phosphate, while maltose and trehalose levels remained unchanged (Figure 5D). Together, these data support a model in which ADP-glucose accumulation contributes to growth toxicity during co-catabolism of glycerol and sugar, consistent with ADP-glucose-mediated toxicity described in other systems^40^. To investigate the impact of Δ*mak glgC*::IS*6110* on antibiotic persistence, we performed a PEM killing assay by treating Δ*mak glgC*::IS*6110* with a combination of INH+RIF. This revealed enhanced persistence (by ∼1000-fold) in the Δ*mak glgC*::IS*6110* strain compared to Δ*mak*, implicating ADP-glucose accumulation in the persistence defect observed in Δ*mak* (Supplemental Figure 7).

### Maltokinase supports *Mtb* survival during persistent infection *in vivo*

Finally, we sought to determine whether the Δ*mak* phenotype extends *in vivo*. C57BL/6 mice were aerosol-infected with a low dose of wild-type, Δ*mak*, or the complemented strain. Δ*mak* established infection at levels comparable to wild-type but exhibited significant attenuation in lungs and spleens at six and eight weeks post infection (Figure 6), indicating a defect in maintenance of infection rather than in initial establishment. Complementation partially restored lung and spleen burden, suggesting that proper regulation or expression context of *mak* contributes to *in vivo* fitness.

**Figure 6.**
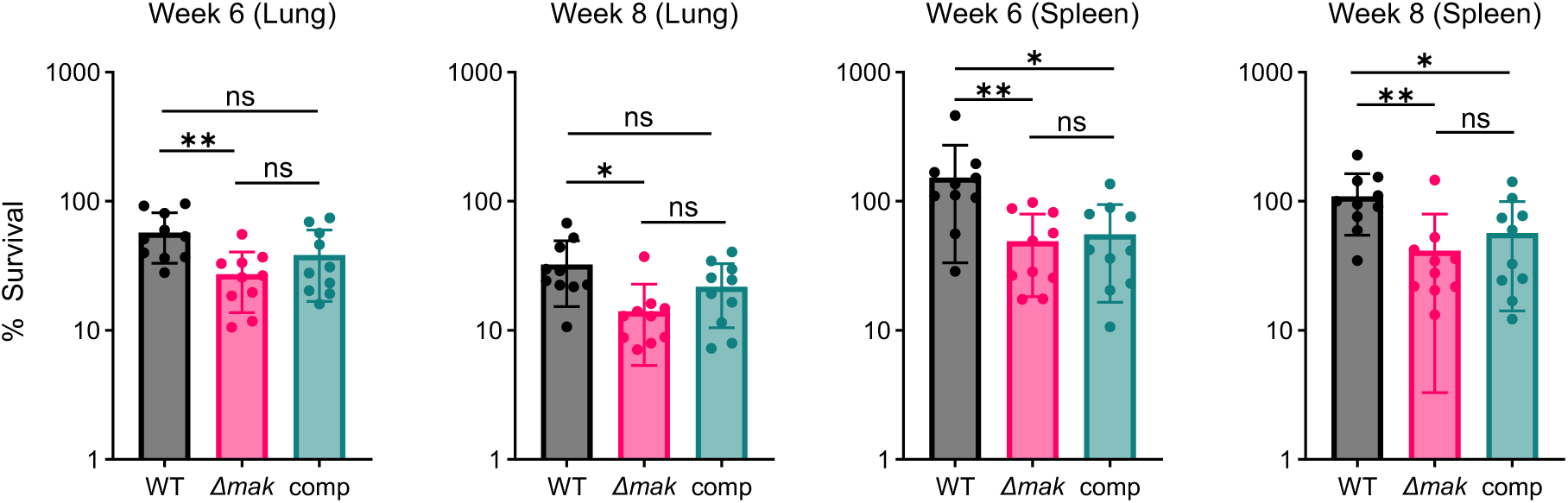
Δ*mak* exhibits attenuation during persistent phase of infection *in vivo.* C57BL/6 mice were aerosol-infected with H37Rv (black), H37Rv Δ*mak* (pink), and the complemented strain (green). Lungs and spleens were harvested at 3, 6, and 8 weeks post-infection. CFU values were normalized to bacterial abundance at 3 weeks post-infection. Data represent mean ± SD from two independent experiments (n = 10). Statistical significance was determined by one-way ANOVA with Tukey’s multiple comparisons test at each timepoint within each organ. (*p <0.05, **p <0.01, ***p <0.001, ****p <0.0001).

Under INH+RIF treatment, Δ*mak* showed a modest reduction in spleen burden and no detectable difference in lungs relative to wild-type (Supplemental Figure 8). The absence of a treatment-specific effect may reflect confounding by baseline attenuation and organ-specific differences in drug exposure and physiology, including the presence of glycerol in mouse lungs^33^. Together, these data indicate that *mak* contributes to *in vivo* fitness during the persistent phase of infection.

## Discussion

Combination chemotherapy has been central to TB treatment since the introduction of isoniazid-containing regimens, yet the cellular mechanisms that enable *Mtb* to survive bactericidal drug combinations remain incompletely understood^41^. Here, we establish a rapid and experimentally tractable Persister Enrichment Model (PEM) that increases the frequency of multidrug-tolerant survivors and enables direct interrogation of the transition from exponential growth to sustained drug tolerance. Using PEM, we show that the temporal structure of drug exposure shapes multidrug tolerance *in vitro* and *in vivo* and identify maltokinase as a metabolic determinant of multidrug persistence *in vitro* and bacterial survival during persistent infection *in vivo*.

PEM differs from commonly used starvation, stationary phase, and hypoxia models in that bacteria are actively replicating immediately prior to antibiotic exposure, allowing examination of early events that commit populations to a tolerant state^19,20,42^. PEM is also rapidly implemented and yields a stable tolerant population that can be exploited for multidrug susceptibility testing. Even low concentrations of glycerol abolish persister enrichment, whereas high dextrose fails to suppress it, arguing against simple carbon starvation as the cause of tolerance. These observations support a model in which glycerol-associated physiology antagonizes entry into multidrug tolerance. Together, these features make PEM a useful complement to existing persistence models, particularly for studying multidrug tolerance during combination therapy.

A major advantage of PEM is the ability to study isoniazid persisters over extended time frames without rapid takeover by resistant mutants. Using this system, we show that INH-tolerant populations display heterogeneous cross-tolerance and importantly, that drug order influences killing by the INH-RIF combination. Simultaneous INH+RIF exposure induces a more robust tolerant population than staggered treatment, and initiating RIF prior to INH reduces the efficacy of the combination both *in vitro* and *in vivo*. These findings provide experimental support for the concept that multidrug tolerance is reactive and history-dependent, as previously proposed^5^. Although TB drugs are administered concurrently in clinical practice, differences in pharmacokinetics and lesion penetration are likely to impose effective temporal ordering *in vivo*, suggesting that drug scheduling may influence the likelihood of entering multidrug tolerance.

Using Tn-Seq under multidrug pressure, we identified *mak*, the maltokinase-encoding gene, as the most strongly depleted gene among our significant hits in survivors formed in PEM. Targeted deletion of *mak* sensitized *Mtb* to INH+RIF *in vitro* and resulted in attenuation during the persistent phase of infection *in vivo*. These findings place maltokinase within a growing set of metabolic factors linked to persistence, including TreS and enzymes of the α-glucan pathway^26,27,29,43–45^.

Our metabolomic and growth data suggest that maltokinase functions at the interface of sugar utilization and storage, enabling excess carbon to be routed into glycogen and thereby supporting co-catabolism of multiple carbon sources. In Δ*mak*, this buffering capacity is impaired, leading to accumulation of intermediates, such as ADP-glucose, that can lead to toxic effects^40^. Perturbations in glycerol-3-phosphate metabolite pools and related metabolites appear to reflect this disruption in carbon buffering, as these elevations are reduced upon *glgC* disruption. Oleic acid partially alleviates these defects, potentially by enabling sequestration of excess carbon through TAG synthesis, whereas its absence exacerbates growth impairment, consistent with a role for alternative carbon sinks in mitigating co-catabolic stress. The accumulation of maltose and ADP-glucose is consistent with a bottleneck in glycogen synthesis, and the partial rescue of the phenotype by *glgC* disruption supports a direct contribution of ADP-glucose buildup to both growth toxicity and the reduced capacity to enter multidrug tolerance in line with observations in other systems^40^. Together, these data support a model in which maltokinase maintains carbon distribution balance during co-catabolism of multiple carbon sources. In its absence, simultaneous carbon influx from glycerol and sugar cannot be efficiently coordinated, leading to accumulation of toxic intermediates and subsequent impairment of growth and persistence.

The glycerol dependence of PEM highlights the broader importance of carbon source context in shaping drug responses. Glycerol availability modulates antibiotic efficacy^23^ and host susceptibility to TB^46^, and reversible variation in the glycerol kinase gene, *glpK*, has been linked to transient drug tolerance^33^. Given *Mtb*’s capacity for co-catabolism of multiple carbon sources^32,47^ we propose that robust buffering of carbon distribution contributes to stable multidrug tolerance, and that perturbation of buffering nodes such as maltokinase increases metabolic liability during antibiotic stress. Although Δ*mak* is attenuated *in vivo*, baseline fitness defects likely complicate detection of treatment-specific persistence phenotypes, particularly in the lungs. Furthermore, given the apparent glycerol-dependency of the persistence phenotype in Δ*mak*, it is possible that glycerol present in the lungs of mice obviates the instability introduced by Δ*mak.* We hypothesize that the survival advantage provided by *glpK* mutants would be attenuated by the simultaneous deletion of *mak*.

In summary, PEM provides a rapid platform to study multidrug tolerance in *Mtb*, reveals that the order and timing of drug administration impact the emergence of multidrug-tolerant populations, and identifies maltokinase as a metabolic determinant of multidrug persistence. This work further explores the TreS-Mak-GlgE axis, providing more context for the interpretation of previous work^27,29,45^. The findings in this study highlight carbon distribution buffering pathways as promising targets for strategies to weaken multidrug-tolerant populations and improve the efficacy of combination TB therapy.

## Materials and methods

### Bacterial strains and culture conditions

Bacterial strains were obtained from laboratory stocks. *Mtb* mc^2^6230 (H37Rv Δ*RD1* Δ*panCD*)^48^ was used as the parental strain for mutant generation, killing assays, growth experiments, and Tn-Seq unless otherwise noted (Supplemental Table 6). *Mtb* H37Rv was used for BSL-3 kill curves and mouse infection experiments (Supplemental Table 6). Bacteria were grown in Middlebrook 7H9 liquid medium (BD Difco) supplemented with 0.05% Tyloxapol (Sigma-Aldrich), 0.1 mM sodium propionate (Sigma-Aldrich), and OADC (final 0.06% sodium oleate (Strem Chemicals), 5% albumin (GoldBio), 2% dextrose (Fisher Chemical), 0.03% catalase (Sigma-Aldrich), and 0.085% NaCl) +/- 0.5% glycerol (Fisher Chemical). This medium is referred to as Dex2 with or without glycerol. *Mtb* mc^2^6230 cultures were additionally supplemented with 24 mg/L pantothenate. Media for high-dextrose conditions was additionally supplemented with dextrose to a final concentration of 50 mg/mL, deemed Dex50 media. 40 μg/mL kanamycin or 75 μg/mL hygromycin were used when required for antibiotic selection.

Middlebrook 7H10 solid agar (BD Difco) supplemented with 0.5% glycerol, 1 mM sodium propionate, 0.06% sodium oleate, 5% albumin, 2% dextrose, 0.03% catalase, and 0.085% NaCl was used for agar plates. 40 μg/mL kanamycin or 75 μg/mL hygromycin were used when required for antibiotic selection. Plates used with BSL-2 strains were additionally supplemented with 24 mg/L pantothenate.

For all experiments, starter cultures were inoculated from frozen seed stocks and subcultured once before use in experiments. Subcultures were generally grown for 4 days to an optical density OD_600_ of ∼0.8. Subcultures for persister enrichment experiments were grown in the absence of glycerol for the same amount of time as glycerol-containing cultures, typically to an OD_600_ of ∼0.4.

### Growth assays

Strains were precultured in Dex2 media with glycerol to OD_600_ of ∼0.8, subcultured into fresh Dex2 media with glycerol at a calculated starting OD_600_ of 0.025, and grown for 4 days to OD_600_ of ∼0.8. Cultures were then washed three times in carbon-free 7H9 medium supplemented with albumin, catalase, sodium chloride, and 0.05% tyloxapol to remove residual carbon sources before inoculation into inkwells containing experimental media at a calculated starting OD_600_ of 0.0033.

### Killing assays

Strains were precultured in Dex2 media with glycerol to an OD_600_ of ∼0.8. Cells were then washed three times in Dex2 media, and then sub-cultures were inoculated at a calculated OD_600_ of 0.025 in either Dex2 with or without glycerol. Cultures from Figure 1F were sub-cultured in carbon matched Dex8 (8 mg/mL dextrose) or low glycerol (0.162%) media where appropriate. After 4 days, cells grew to an OD_600_ of ∼0.8 for glycerol cultures and ∼0.4 for non-glycerol cultures. Cells were then diluted to a calculated OD_600_ of 0.33 (10^8^ CFU/mL), and then drugs prepared in either water or DMSO were added at 50-100× MIC concentrations. Drug concentrations were INH (36 µM), RIF (3 µM), EMB (490 µM), LNZ (88 µM), OFX (138 µM), and DCS (980 µM). At each timepoint, cells were sampled and plated on standard 7H10 agar plates. Phosphate-buffered saline +tyloxapol 0.05% was used for serial dilution. Colonies were counted after 3-4 weeks.

### Mutant generation and complementation

Deletion of *mak* was carried out by specialized transduction as previously described^28^. *Mtb* H37Rv AE1001^49^, a PDIM+ isolate of H37Rv, was used to generate BSL-3 mutants, while *Mtb* mc^2^6230 AE1601, a PDIM+ isolate of mc^2^6230^34^, was used to generate BSL-2 mutants. Confirmation of gene deletion was done by 3-primer PCR and whole-genome sequencing (WGS) as below.

Complementation was carried out using the native promoter of *mak* (250 bp upstream) in a pMV306 plasmid with the L5 integrase removed, alongside a suicide plasmid pYUB3003 containing the L5 integrase to integrate the complement at the L5 insertion site, using a previously described approach^50^.

Oligonucleotides used in this study are listed in Supplemental Table 7. Plasmids were constructed using Gibson assembly using the NEBuilder HiFi DNA Assembly Cloning Kit (New England Biolabs).

*Mtb* mc^2^6230 Δ*mak glgC::*IS*6110* and *Mtb* mc^2^6230 Δ*mak sugI* G62D suppressor mutants were isolated by picking single colonies into standard Dex2 culture media with glycerol, and mutations were identified by WGS. The *glgC*::IS*6110* insertion mutation was additionally confirmed by Sanger sequencing, and WGS was used to ensure the strains did not carry additional SNPs (Supplemental Table 4).

### Whole genome sequencing and analysis

Genomic DNA was extracted and isolated using a CTAB extraction method^51^. DNA was sequenced by Illumina NextSeq performed by SeqCenter (Pittsburg, PA), using the Illumina DNA Prep kit and sequences on an Illumina NextSeq 2000 (2 x 151 bp reads). Reads were mapped to the H37Rv NC_000962.3 reference genome, and variants were called using Geneious 2025.0.1. IS*6110* insertions in *Mtb* mc^2^6230 Δ*mak glgC::*IS*6110* colonies were identified using the variant caller Pilon as previously described^34^, as the Geneious variant caller failed to detect mutations in these strains.

### Transposon sequencing (Tn-Seq) and statistical analysis

Preparation, isolation, and DNA processing of a saturated transposon mutagenesis pool was carried out by Himar1 mutagenesis based on a previously published protocol^52^ with a few modifications. To obtain a high-titer lysate of phAE180, single plaques were picked and assessed for mutant formation in *Mycobacterium smegmatis*. Single plaques that yielded the most highly kanamycin-resistant colonies were selected, and single plaques were picked again. The plaque yielding the most kanamycin-resistant colonies was amplified on agar plates and purified by CsCl_2_ ultracentrifugation. To make saturated transposon mutagenesis pools, 300 mL of *Mtb* at OD_600_ ∼0.8 was infected with 3 mL of high-titer lysate (MOI of 1), then plated on large 25×25 cm agar plates (12 plates total). The resulting colonies from these plates were scraped and combined into Dex2 media with glycerol. The pool was plated to enumerate CFUs, then cultures were inoculated at a calculated 3×10^6^ CFU/mL in 50 mL of media in roller bottles to preserve the complexity of the transposon mutagenesis pool. This pool was grown to ∼1×10^8^ CFU/mL (∼4 days), before being treated with 50-100× MIC drug concentrations for 7 days. A calculated 3×10^6^ CFU were sampled from these pools after drug treatment, based on the number of surviving bacteria observed in previous experiments, washed to remove residual drugs, and regrown in fresh media for 6 days to enrich for surviving mutants. DNA was extracted from these cultures and then processed according to established Tn-Seq sample preparation protocols^52,53^ with alterations as below.

At the step of amplifying transposon-gene junctions, Q5 polymerase was used in place of Taq, with 20 amplification cycles using the following buffer/PCR parameters: Q5 reaction buffer: 40 μL (0.5 final), Q5 GC enhancer: 40 μL (0.5 final), dNTPs (10 μM): 8 μL, Primer1: jel_ap1 (10 μM): 20 μL, Primer2: T7 (10 μM): 20 μL, DMSO: 20 μL, Q5 polymerase: 4 μL, DNA (160 ng, 20 ng per reaction), water to 400 μL. The reaction was then split over eight tubes (50 μL per reaction). PCR thermal cycling conditions were 98 °C for 30 s, followed by 20 cycles of 98 °C for 10s, 65 °C for 20 s, 72 °C for 45s, with a final extension at 72 °C for 2 min.

After DNA was isolated by gel extraction, sequencing primers were applied to the libraries using Q5 polymerase with 10 amplification cycles using the following buffer/PCR parameters: Q5 reaction buffer: 4 μL, Q5 GC enhancer: 2 μL, dNTPs (10 μM): 0.625 μL, Primer1: Solmar (1 μM): 2 μL, Primer2: Sol_AP_XXX (1 μM): 2 μL, Q5 polymerase: 0.2 μL, DNA 20 ng, water to 20 μL. PCR thermal cycling conditions were 95 °C for 30 s, followed by 10 cycles of 95 °C for 30s, 58 °C for 30s, 72 °C for 45s, with a final extension at 72 °C for 5 min. The resulting DNA products were purified via AMPure beads (Beckman Coulter, USA). Other manipulations of DNA were carried out in accordance with the referenced protocol^52^.

Sequencing was performed using Nextseq 500 (2 × 150 bp paired-end reads). The sequenced reads were demultiplexed and processed using TPP (Transit Pre-Processor), a pre-processor included in the TRANSIT software^54^. Statistical analysis was performed using a resampling analysis in TRANSIT with a beta-geometric correction^55^. Samples treated in the presence of glycerol were treated as the control strain and those treated in the absence of glycerol were used as the treatment strain within the TRANSIT software. The *Mtb* H37Rv reference genome (NC_000962.3) was used to map transposon-chromosome junctions using the Burroughs Wheeler Aligner in TRANSIT^56^.

Processed Tn-seq data from Figure 3B can be found in Supplemental Table 3.

### Metabolite extraction

Triplicate roller bottles containing 50-100 mL of standard 7H9 media with various carbon sources (Dex2 +0.5% glycerol, Dex50 +0.5% glycerol) were inoculated at a starting OD_600_ of 0.05, and then metabolites were extracted after 5 days of growth. An equivalent of 10 mL of culture at an OD_600_ of 0.2 was rapidly filtered on 0.45 μm Durapore PVDF membrane filters (MilliporeSigma) using a vacuum manifold (MilliporeSigma). Cultures were quenched by placing the filter paper in 1 mL of extraction solvent containing 20:40:40 (v/v) water/acetonitrile/methanol with approximately 500 μL of 0.1 mm zirconia/silica beads (BioSpec) at −20 °C. Samples were homogenized using a Precellys Cryolys Evolution (Bertin Technologies) cooled to 0 °C for three 20 s cycles at 6800 rpm with a 30 s pause between cycles. Samples were centrifuged, and the extracts were filtered through a 0.22 μm Nylon Spin-X microcentrifuge filter (Corning) and stored at −80 °C. For analysis, extracts were concentrated 5-fold using a SpeedVac^®^ Plus SC110A (Savant Instruments, Inc.) to evaporate the solvent, and then redissolved in 1/5^th^ volume of the extraction solvent.

### LC-MS metabolomic analysis

Metabolomics analysis was performed using an Agilent 1290 Infinity II liquid chromatography (LC) system coupled with an Agilent 6545 quadrupole time-of-flight (QTOF) mass spectrometer (MS) equipped with a Dual Agilent Jet Stream Electrospray Ionization (Dual AJS ESI) source operated in negative mode as previously described^34^. Data analysis was performed using the Agilent MassHunter Qualitative and Quantitative Analysis Software. Metabolite identification was based on mass-retention times determined using chemical standards. For semi-targeted analysis, the Sigma MSMLS metabolite reference library (Sigma-Aldrich) was used for peak annotation as previously described^35^. Metabolites were quantified using a mass tolerance of 20 ppm with manual curation of peak areas where necessary, and the area under the curve (AUC) was determined.

### Mouse experiments

Mouse experiments were performed in accordance with National Institutes of Health guidelines following the recommendations in the Guide for the Care and Use of Laboratory Animals. The protocol used in this study was approved by the Institutional Animal Care and Use Committee of Albert Einstein College of Medicine (Protocol #00001445). Female C57BL/6 mice (Jackson Laboratory) were infected with H37Rv via the aerosol route using a 1×10^7^ CFU/mL *Mtb* suspension in PBS containing 0.05% tyloxapol and 0.004% antifoam. Mice were sacrificed and the right lung and spleen were homogenized in PBS and then serial dilutions plated on 7H10/OADC/glycerol plates at each timepoint to determine CFU per organ. Plates were incubated for 3 weeks before counting colonies. Mice with no detectable CFUs and exhibiting greater than a 2-log difference relative to the other replicate mice were excluded from analysis.

## Acknowledgements

We thank Bing Chen and John Kim for assistance with animal experiments; Saranathan Rajagopalan for the pYUB3003 plasmid, and the laboratory of William R. Jacobs Jr for their gifts of phAE180 and the transducing phages; Thomas R. Ioerger, Anthony D. Baughn, Christopher Sassetti, and Michael DeJesus for their guidance and assistance with Tn-Seq methodology and interpretation. We thank the Albert Einstein College of Medicine epigenomics core for assistance in sequencing Tn-Seq libraries. Graphical illustrations were created with BioRender.com. Generative AI tools (ChatGPT, OpenAI) were used for limited editorial assistance, including grammar review, wording suggestions, and formatting feedback. All scientific content, interpretation, analysis, and final writing decisions were performed by the authors. M.W.S acknowledges support from the Institutional AIDS training grant, Training in HIV/AIDS Pathogenesis; Basic and Translational Research (T32 AI007501), and the Albert Einstein College of Medicine MSTP training grant (T32 GM149364). C.V.M., M.C., J.C., and M.B. acknowledge support from the National Institutes of Health/National Institute of Allergy and Infectious Diseases (R01 AI139465 and R01 AI175972), the Potts Memorial Foundation, and Albert Einstein College of Medicine internal funding.

## Data availability

Whole genome sequence data and raw transposon mutagenesis sequence data have been deposited in the NCBI Sequence Read Archive (SRA) under the Bioproject accession number PRJNAXXXXXX.

## Supplemental Material

Supplemental Figures and Tables. Figures S1-S8, Tables S1, S4-7

Supplemental Table 2.

Supplemental Table 3.

